# AlphaFold 2, but not AlphaFold 3, predicts confident but unrealistic β-solenoid structures for repeat proteins

**DOI:** 10.1101/2024.10.30.621056

**Authors:** Olivia S. Pratt, Luc G. Elliott, Margaux Haon, Shahram Mesdaghi, Rebecca M. Price, Adam J. Simpkin, Daniel J. Rigden

## Abstract

AlphaFold 2 has revolutionised protein structure prediction but, like any new tool, its performance on specific classes of targets, especially those potentially under- represented in its training data, merits attention. Prompted by a highly confident prediction for a biologically meaningless, scrambled repeat sequence, we assessed AF2 performance on sequences comprised perfect repeats of random sequences of different lengths. AF2 frequently folds such sequences into β-solenoids which, while ascribed high confidence, contain unusual and implausible features such as internally stacked and uncompensated charged residues. A number of sequences confidently predicted as β-solenoids are predicted by other advanced methods as intrinsically disordered. The instability of some predictions is demonstrated by Molecular Dynamics. Importantly, other Deep Learning-based structure prediction tools predict different structures or β-solenoids with much lower confidence suggesting that AF2 alone has an unreasonable tendency to predict confident but unrealistic β-solenoids for perfect repeat sequences. The potential implications for structure prediction of natural (near-)perfect sequence repeat proteins are also explored.

## Introduction

Accurate protein structure determination is essential for understanding biological processes and protein-related disease. Protein structures are experimentally determined mainly by protein crystallography or cryogenic electron microscopy (cryoEM) but these approaches can be very time-consuming, with single structures sometimes taking months or even years to solve (McPherson & Gavira, 2014). For this reason, there has been great interest in methods for computational structure prediction, especially the ab initio of de novo methods that are independent of the availability of already known similar structures (templates). In 2020, at the 14th Critical Assessment of Protein Structure Prediction (CASP14), AlphaFold2 (AF2), a deep-learning based model, was introduced by Google DeepMind (Moult *et al*., 2020) and was recognised as a huge step forward in prediction accuracy in the absence of templates (Jumper *et al*., 2021). AF2 was then deployed on a large scale resulting in the AlphaFold Protein Structure Database (AFDB) (Varadi *et al*., 2024) which has over 214 million models.

Like previous generations of modelling software (Marks *et al*., 2012), accurate template-independent structure prediction with AF2 depends (Jumper *et al*., 2021) on the availability of a sufficiently large and diverse multiple sequence alignment (MSA). Analysis of the MSA highlights amino acid covariance i.e., which pairs of residues have co-evolved and are therefore likely to be close in 3D space in the folded structure. In addition to coordinates, AF2 produces two quality estimates. The pLDDT (predicted Local Distance Difference Test; Mariani *et al*., 2013) is a per-residue local structure confidence score, ranging from 0-100, which is returned in the B-factor column of the structure. By convention, pLDDT is coloured from blue (high confidence) to red or orange (low confidence). The second measure, the PAE (Predicted Align Error), informs on global structural prediction confidence e.g., whether inter-domain orientations can be trusted (Jumper *et al*., 2021).

AF2 can be utilised to explore understudied proteins and plug gaps in our protein structural knowledge. Here we apply AF2 to repeat proteins, which possess sections of repetitive sequences called tandem repeats. These repeats are very diverse, from the repetition of a single residue to hundreds (Kajava, 2012). Despite occurring in ∼14% of proteins (Marcotte *et al*., 1999), they remain poorly characterised (Luo & Njveen, 2014). To address this, in 2014 the database RepeatsDB was launched to automate the annotation and classification of repetitive structures (Domenico *et al*., 2014). It recognises five classes (Kajava, 2012).

Class I consists of crystalline aggregates which have short repeat units between 1-2 residues, often associated with neurodegenerative disorders (Karlin *et al*., 2002).

Class II refers to fibrous structures with repeats between 3-4 residues, for example, collagen. The most researched class, class III, are elongated structures which have repeats between 5-40 residues, consisting of solenoid and non-solenoid proteins.

Class IV are ‘closed’ structures that have repeat lengths overlapping with classes III and V, but fold circularly. Finally, class V or ‘beads on a string’ structures can fold into independent stable domains, such as the DNA binding Zinc-finger domain (Lee *et al*., 1989).

Recently there has been increased interest in beta-solenoid (β-solenoid) proteins within class III. With repeat units between 5-30 residues, β-solenoids display repeating parallel beta strands separated by tight turns (Kajava & Steven, 2006), but due to their vast architectural diversity, they remain largely uncharacterised. In 2023, AF2 was used to predict the structures of several new β-solenoids (Mesdaghi *et al*., 2023), highlighting disease-related repeats such as functional amyloids and the *Chlamydia trachomatis* protein PmpD.

As mentioned, for proteins that lack templates in the Protein Data Bank (Berman *et al*., 2003), AF2 requires a sufficiently information-rich MSA for accurate structure prediction. Thus, for singleton proteins that lack homologues, AF2 typically struggles to make confident predictions. However, surprisingly, we observed that when the repetitive sequence of human protein mucin 22 is scrambled, AF2 still predicts a confident [-solenoid. Furthermore, the model appears implausible, showing uncompensated internal stacking of glutamic acid, which is unrealistic for natural and stable protein structures due to repulsion of the negative charges (Figure 1).

**Figure 1:**
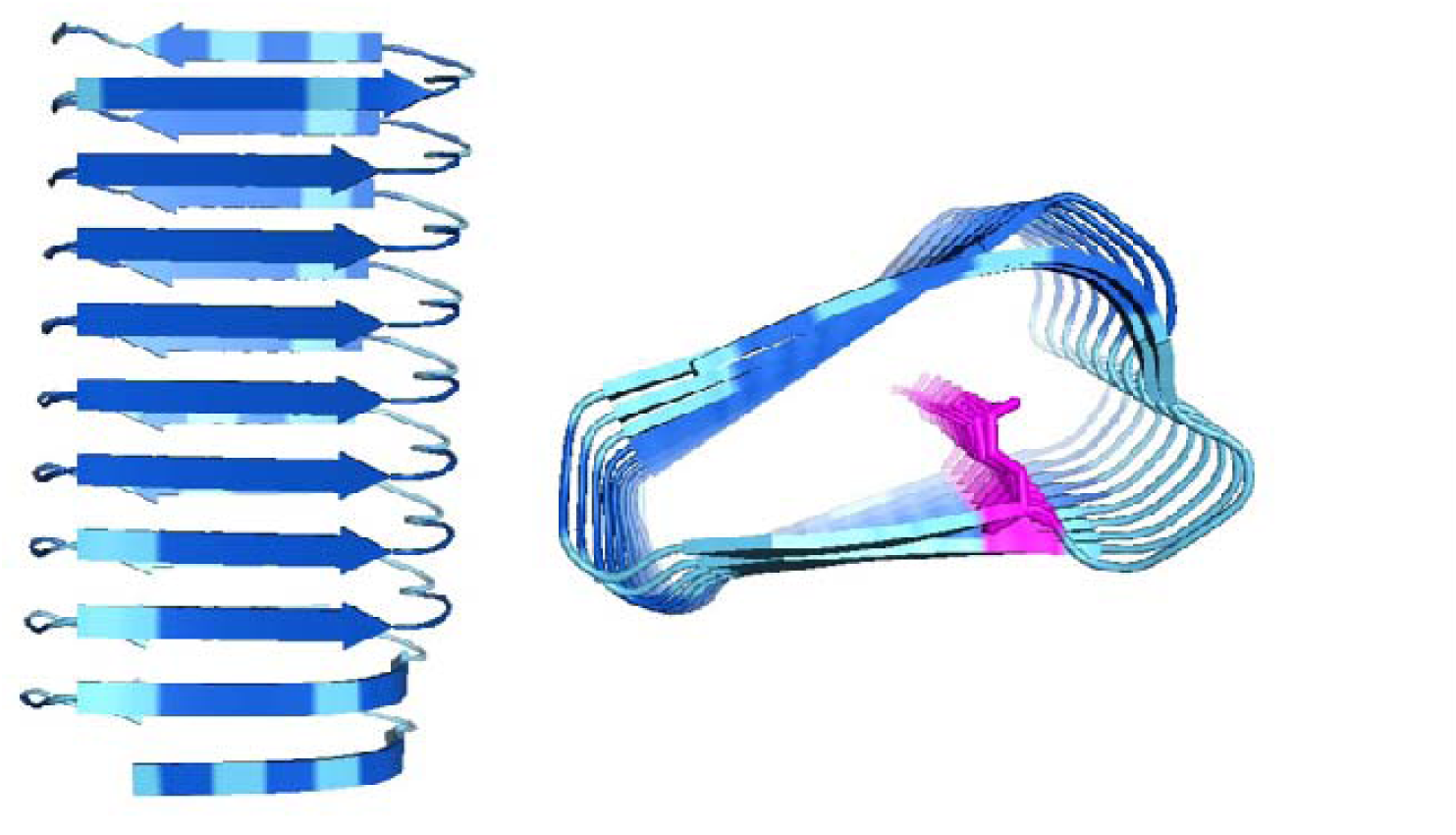
AF2 model from scrambling the repetitive sequence of protein mucin 22. The sequence was randomised into a 20-residue unit (sequence QTSTVIGTIASTETSSSTGI), and concatenated 10 times to form a 200-residue repeat protein comprised of 10 rungs. There is an uncompensated internal glutamic acid stack (pink), which is generally unfavourable to be within the protein core. Despite this, the model is highly confident (pLDDT 90.3) Like later figures, the model backbone is coloured by pLDDT: Dark blue for pLDDT>90, cyan for 70 < pLDDT <90, yellow for 50 < pLDDT <70, orange for pLDDT <50. Figure made using PyMOL (pymol.org).

Consequently, we hypothesised that AF2 may have a bias towards predicting [- solenoids, which could affect structure prediction of natural repeat proteins, including those involved in disease. This report therefore aims to investigate this potential blind-spot.

## Methods

### Sequence & model generation

Firstly, the ‘RANDOM’ function in Python (v3.11) was used to create random 20- residue sequences, concatenated 10 times, producing 200-residue repeat sequences. The sequences were designed to cover a range of amino acid propensities for alpha and beta secondary structure (Costantini *et al*., 2006) and the average propensity scores for both were calculated by taking the mean for each sequence. 50 sets of models were generated using the AlphaFold2 Colab page (v1.5.5) (Mirdita *et al*., 2022) under default settings.

The same process was then repeated, altering the repeat unit length to be between 5-30 residues. For each length, three random sequences were created: one biased towards beta secondary structure, one towards both alpha and beta, and one completely random. The sequences were then concatenated 10 times. Models were generated by AF2 locally (v1.5.2), using the parameter *--num-models 5*, resulting in a dataset of 78 sets of models.

### Model classification using STRPsearch

To classify the fold of the rank 1 AF2 models, the software STRPsearch was used (Mozaffari *et al*., 2024), and the E-value set between 0.1-10.0 with the *--max-eval* parameter. As this software uses a library, for models that received no classification or multiple, the most appropriate classifier was manually selected by visualising secondary and tertiary structure.

Calculating RMSD between models 1-5

Using the ‘ALIGN’ command in PyMOL (v2.5.2), for each β-solenoid model, model 1 (the number here referring to the rank) was compared to models 2 through 5, and the root mean square deviation (RMSD) was calculated using the parameters *cycles=0, transform=0*.

Model validation with Verify3D

For the following analysis, model refers to rank 1 models only. To investigate the plausibility of the β-solenoid models, they were submitted to the Verify3D server (Eisenberg *et al*., 1997). The average Verify3D score was calculated by taking the mean of the per-residue average scores. A categorical failure is awarded by Verify3D if fewer than 80% of residues within a model score >=0.1 in the 3D/1D profile.

As a standard, β-solenoid crystal structures identified by RepeatsDB were also analysed by Verify3D. First, the sequences were input into RADAR (Heger & Holm, 2000), to identify those that most exemplified ‘perfect’ sequence repetition. For normalisation, the RADAR score was divided by the repeat unit length. Those with a score above 7.0 were selected for comparison.

### Modelling with ESMFold, AlphaFold3 and RoseTTAFold-All-Atom

The sequences were also modelled using the ESMFold Colab page (Lin *et al*., 2023) and the AlphaFold3 (AF3) server (Abramson *et al*., 2024). To compare the outputs, TM-scores between models for the same sequence were calculated (Zhang & Skolnick, 2004; Xu & Zhang, 2010). Average pLDDT values for the AF3 models were determined with ‘GEMMI’ (Wojdyr, 2022) in Python as AF3 gives atomic pLDDT. Finally, the sequences were modelled with RoseTTAFold-All-Atom (RFAA) (Krishna *et al*., 2024) for additional comparison.

### Disorder predictions with AIUPred

The predicted disorder for each sequence was determined with the AIUPred server (Erdős & Dosztányi, 2024). The average probability was calculated by taking the mean of the per residue probabilities.

### All-atom molecular dynamics with GROMACS

For models that had internally stacked charged residues and β-solenoid crystal structure 3JX8:A, all-atom molecular dynamics (MD) was implemented with GROMACS 2022.2 (Van Der Spoel *et al.,* 2005). The models were first solvated with the TIP3P water model (Jorgensen *et al*., 1983) and neutralised with Na^+^/Cl^-^ ions.

Particle mesh Ewald algorithms were used to determine the long-range electrostatic interactions (Pronk *et al*., 2013). Next, energy minimization was carried out for up to 10,000 steps, using the steepest descent followed by the conjugate gradient algorithm. Then, the temperature of the system was raised from 0K to 300K. Following two 20ps equilibration steps, three 1µs simulations were run at a 2 fs timestep. The simulation was implemented with the CHARMM36 force field (Bjelkmar *et al*., 2010) and results visualisation was conducted with VMD 1.9.3 (Humphrey *et al*., 1996). Finally, analysis was performed using the ‘MD-DaVis’ (Maity & Pal, 2022) and ‘MDAnalysis’ (Michaud-Agrawal *et al*., 2011) modules in Python and the RMSD calculated with the ‘rms’ command.

### Sequence alignment with HHPred

Sequences were searched against the Pfam (Mistry *et al.,* 2021) and PDB (Berman *et al*., 2003) databases with HHPred (Zimmermann *et al*., 2018) to search for chance sequence matches.

## Results & Discussion

### Preliminary exploration of 20-residue repeat units highlights potentially problematic predictions

To begin, a small preliminary dataset of β-solenoid models was produced in which each model was of a randomly generated 20-residue repeat unit concatenated ten times. Of the 50 models generated, 36 resulted in a β-solenoid structure but there was some redundancy in the set as some sequences differed only at single positions with others. Nevertheless, these initial models served to identify interesting factors to explore, for example, investigating how model propensity for different secondary structure influences AF2 structure prediction. One particularly concerning observation identified involved the internal stacking of negatively charged residues within the protein core. Additionally, the protein structure validation software Verify3D highlighted several models as implausible, whilst alternative deep-learning modelling methods showed contradictory outputs to AF2. To test these observations on a larger scale, 78 sets of models were generated with AF2, covering a range of repeat unit lengths. The results for this dataset are presented below.

### Expanding the exploration across different repeat unit lengths

AF2 generates confident β-solenoids across different repeat unit lengths Following generation of the larger dataset in which the repeat unit lengths were altered, for each sequence modelled, the average propensity scores for alpha and beta secondary structure were determined and the model (here model refers to the rank 1 AF2 model), was fold classified by STRPsearch (Figure 2). 32 of the 78 models were classified as β-solenoids, with a few having a potential repeat region identified by STRPsearch.

**Figure 2:**
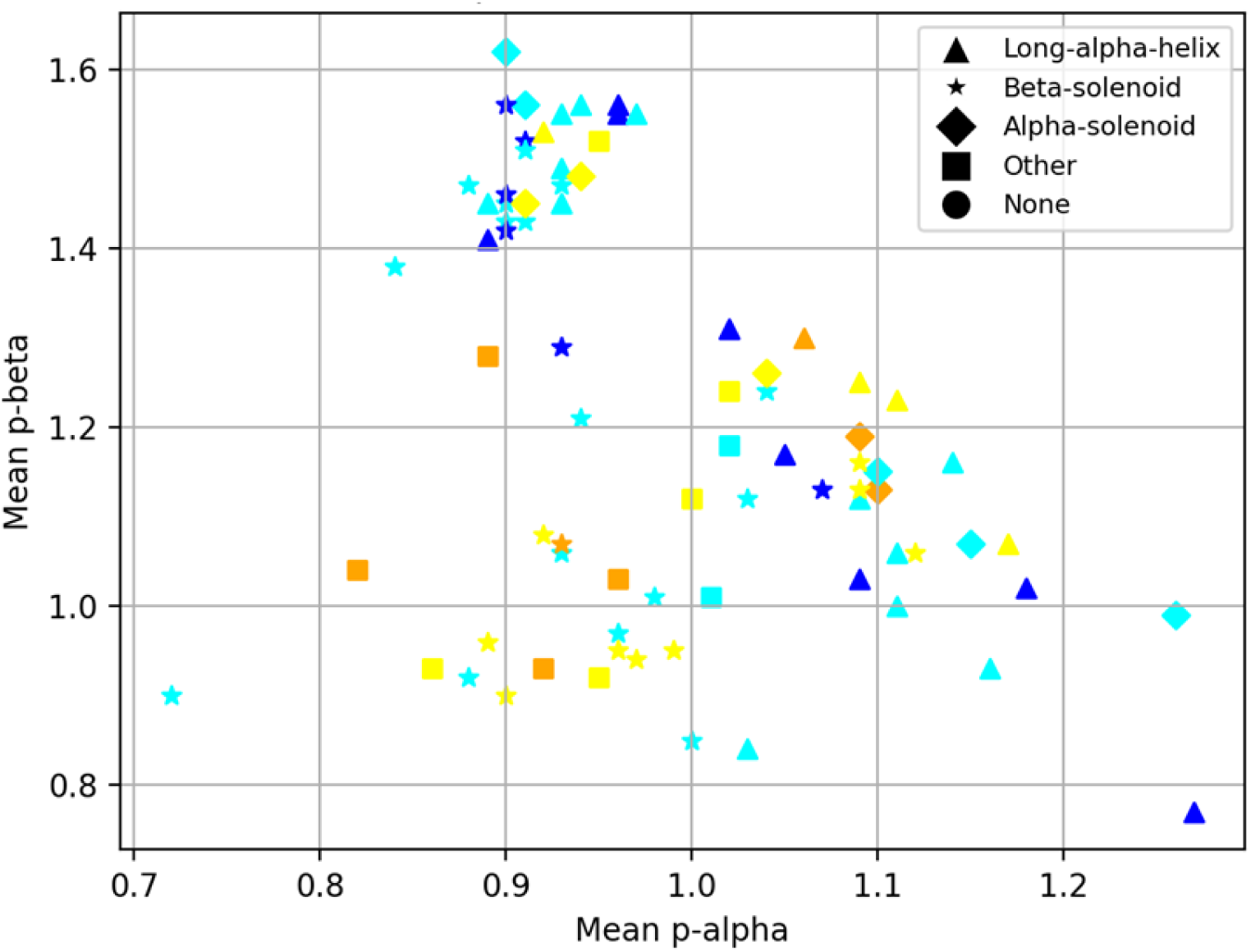
Each point represents a model, its coordinates showing the average sequence propensity score for alpha secondary structure on the x-axis and beta secondary structure on the y-axis. The different markers represent the fold that each model was classified as, coloured by pLDDT - Dark blue for pLDDT>90, cyan for 70 < pLDDT <90, yellow for 50 < pLDDT <70, orange for pLDDT <50. ‘None’ refers to disordered proteins.

Despite being random repeat sequences that lack homologues, AF2 generates many confident β-solenoid models. With regard to secondary structure propensity, it appears that there is no strong pattern, with the exception that above a beta propensity of ∼1.3, there are no low-quality models. Figure S1 highlights that high pLDDT β-solenoids were produced across the size range in 32 of 78 (41%) proteins, suggesting that the potential blind-spot observed in the preliminary exploration may extend across β-solenoids of all lengths.

### Initial model observations show potentially implausible structures

Upon close examination of the β-solenoid models, it appeared that some possessed problematic features. Figure 3a shows a confident model (pLDDT 83.5) with several hydrophobic residues located on the protein surface and obvious clashing of atoms. In addition, Figure 3b displays another confident model (pLDDT 82.4) with aspartic acids stacked internally. Both observations are unlikely to occur within natural and stable proteins. Commonly, hydrophobic residues are found within the protein core, minimising their contact with solvent (Tanford, 1962), and internal negatively charged stacking is energetically unfavourable due to repulsion (Honig & Nicholls, 1995). An exception would be calcium-dependent antifreeze β-solenoid proteins, which have internal negative stacking to coordinate calcium ions (Garnham *et al*., 2011). AF2’s confidence may therefore be explained by the fact that these structures were present in the training data, misleading its predictions. Nevertheless, this does not explain other potentially problematic observations, which reinforce the suggestion that AF2 often produces confident but unrealistic β-solenoid predictions for artificial repetitive sequences.

**Figure 3:**
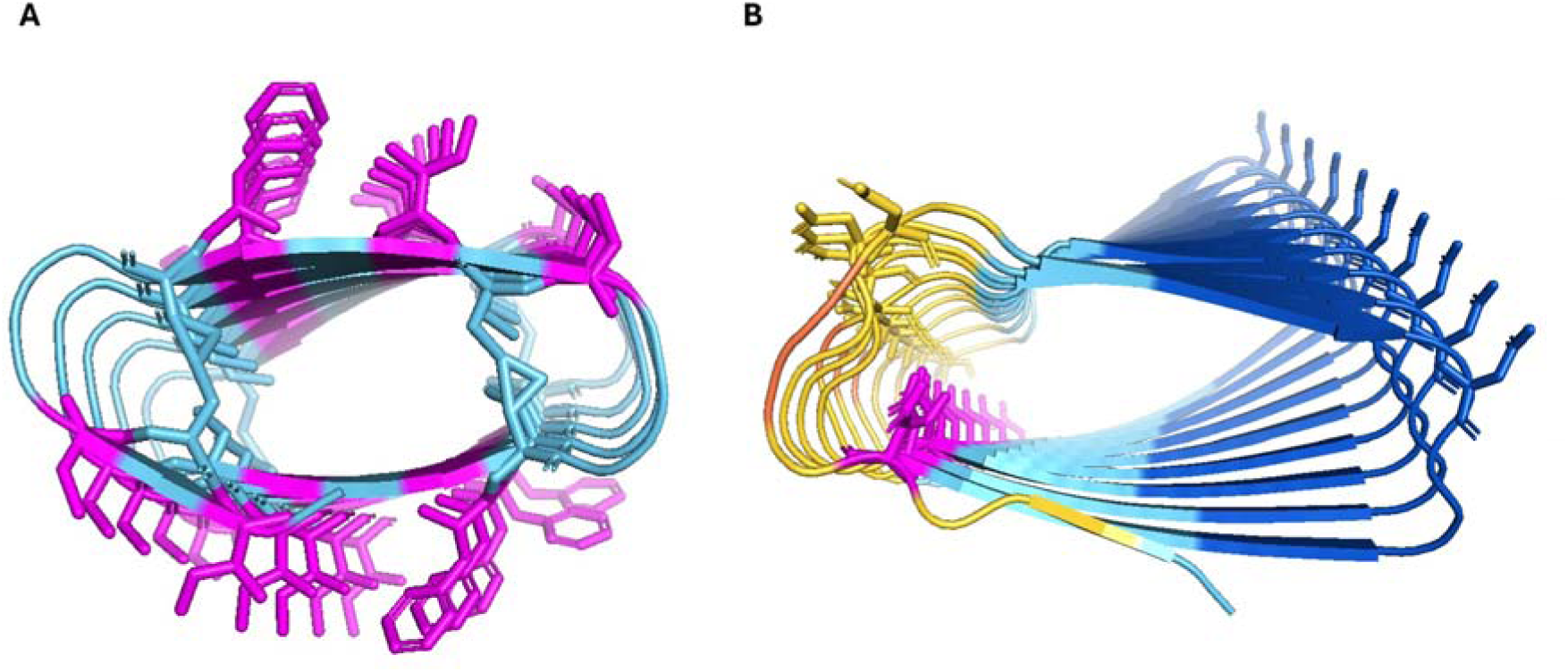
A) A model showing hydrophobic residues (shown as sticks), those on the protein surface are coloured pink. Hydrophobic amino acids are more likely to be found within the protein core. Clashing between residues is also observed. B) A different model showing uncompensated internal stacking of aspartic acid (pink) which is energetically unfavourable due to its negative charge. Figure made using PyMOL (pymol.org).

When analysing the 5 models for each sequence (the number corresponding to the rank), for one interesting case, models 1 and 2 differed significantly in secondary structure. Model 1 displays the β-solenoid fold (pLDDT 91.9), but model 2 appears as a long α-helix (pLDDT 87.2) (Figure 4). Theoretically, both models may be plausible as some proteins are able to adopt multiple conformations (Murzin, 2008) and proteins such as the prion protein (Prp) have both a normal and pathogenic form (Prusiner, 1998), which can differ in secondary structure (Rufai *et al*., 2019). However, these radically different predictions, both confident, more likely further highlight a contradiction in AF2’s predictions.

**Figure 4:**
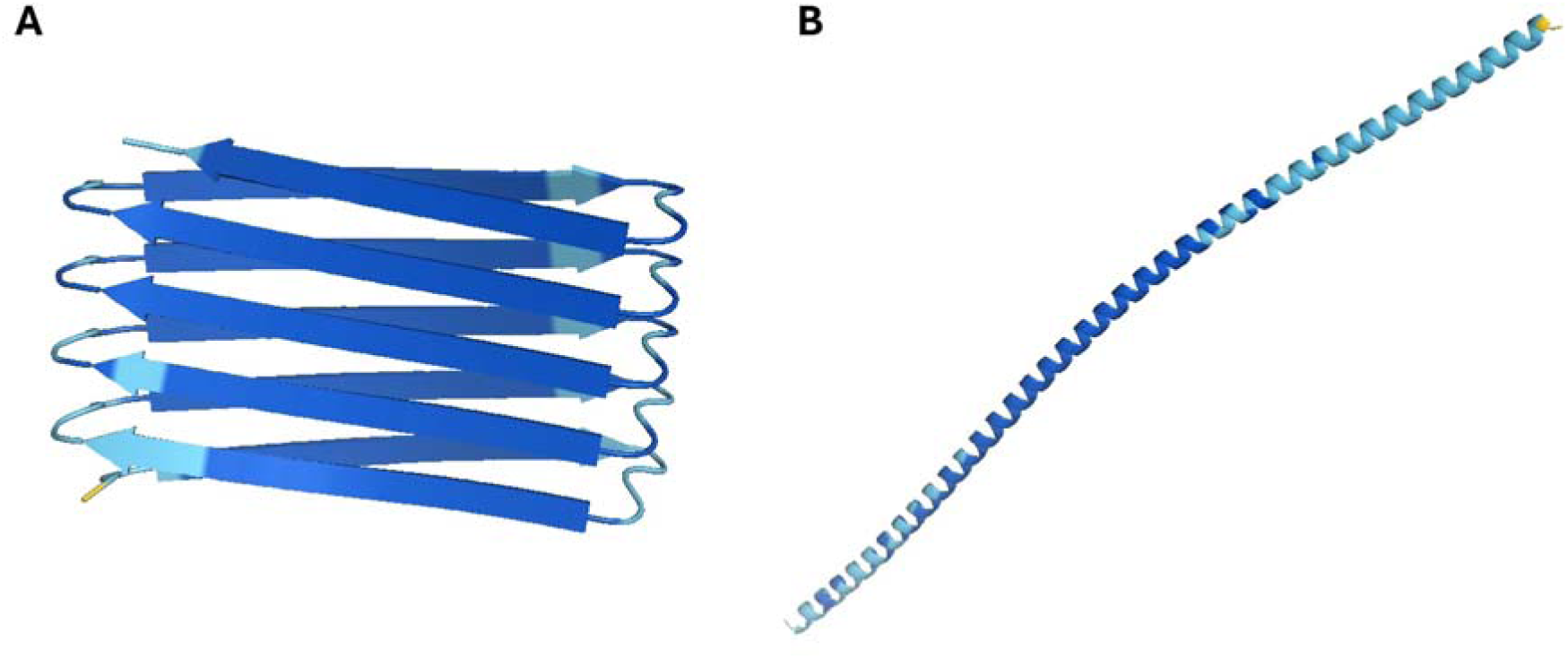
Comparison of AF2 models 1 & 2 from the same random repeat sequence (WCFVCVIVTVFYCW). A) Model 1 (pLDDT 91.9) B) Model 2 (pLDDT 87.2). The RMSD between the two structures is 56.0 Å despite AF2 being confident in both. Figure made using PyMOL (pymol.org).

### Model validation with Verify3D highlights model implausibility

To investigate the plausibility of the β-solenoid models further, they were validated quantitatively through Verify3D. Verify3D considers residue solvent accessibility, secondary structure, and the overall environment to determine whether a protein structure is plausible (Eisenberg *et al*., 1997). To establish the threshold for valid protein structures, seven β-solenoid crystal structures, chosen for the near-perfection of their sequence repeats, were first analysed. For three structures, Verify3D gave a categorical fail, possibly due to the much smaller number of β-solenoid structures available in the PDB when the software was developed. Nevertheless, all seven crystal structures received an average score >0.1, allowing this to be adopted as threshold for interpretation of models.

After analysing the β-solenoid models, 20 out of 32 received an average score <0.1 (Figure 5), 16 of which had a pLDDT >70. This highlights that AF2 generates confident β-solenoid models despite their scoring poorly by validation tools. It was hypothesised that models with a potential repeat region identified by STRPsearch may be more similar to natural proteins, as the software uses a library, and therefore receive a higher Verify3D score. However, no trend is observed.

**Figure 5:**
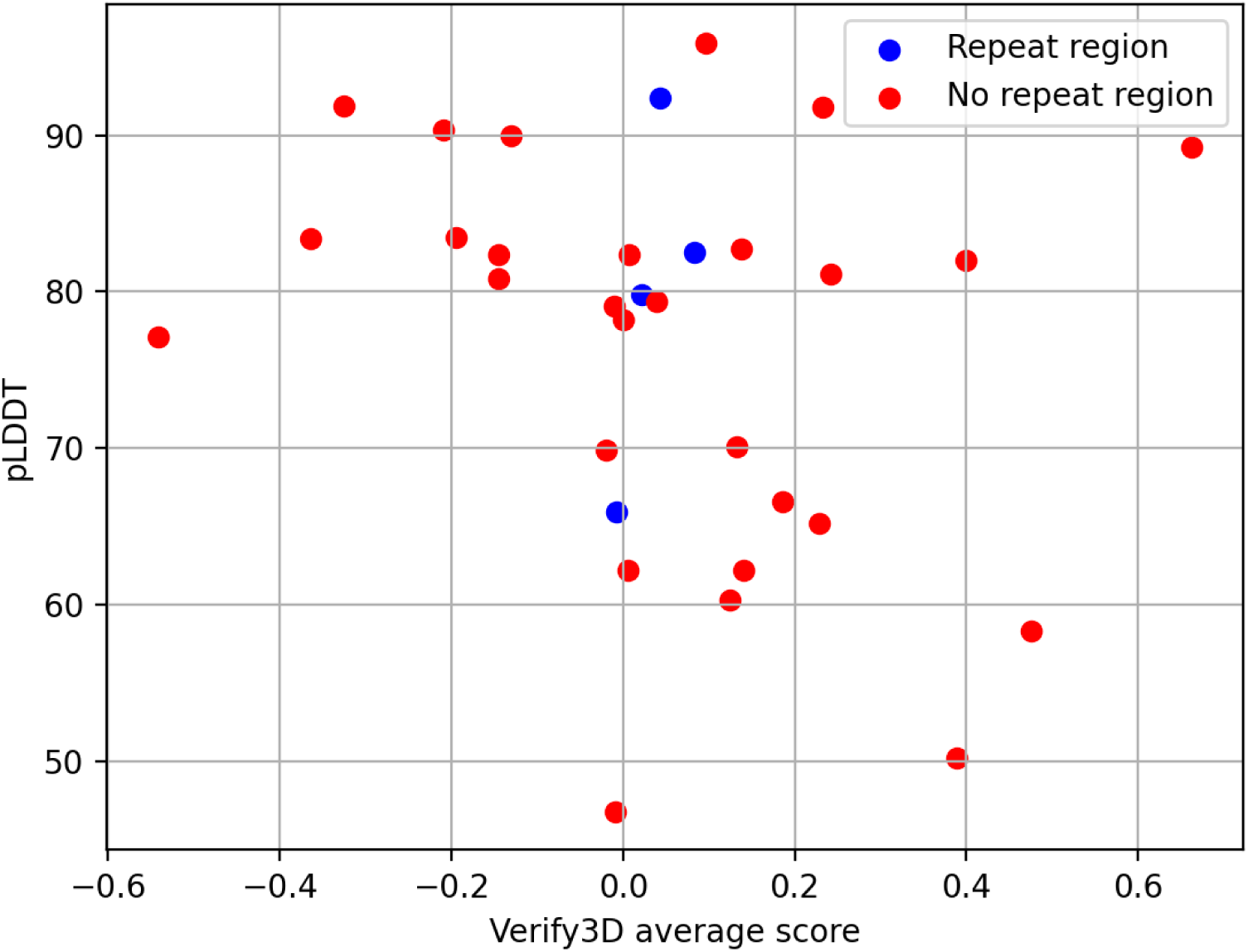
Verify3D results showing the average score for each model on the x-axis against average pLDDT on the y-axis. Each model is coloured by whether a repeat region was identified during classification via STRPsearch (blue for a positive identification, red otherwise).

As the Verify3D score for each model is an average, it was considered possible that problematic features may be neutralised by more favourable ones. For instance, two models received an average Verify3D score >0.1, despite having charged residues stacked internally with high pLDDT. In one model, two internal histidine stacks (Figure 6a), and in the other, several internal charged stacks (Figure 6b) would likely cause significant repulsion. Consequently, AF2 models with a pLDDT >70, but a Verify3D score <0.1 or internal charged stacking, were defined as ‘problematic’.

**Figure 6:**
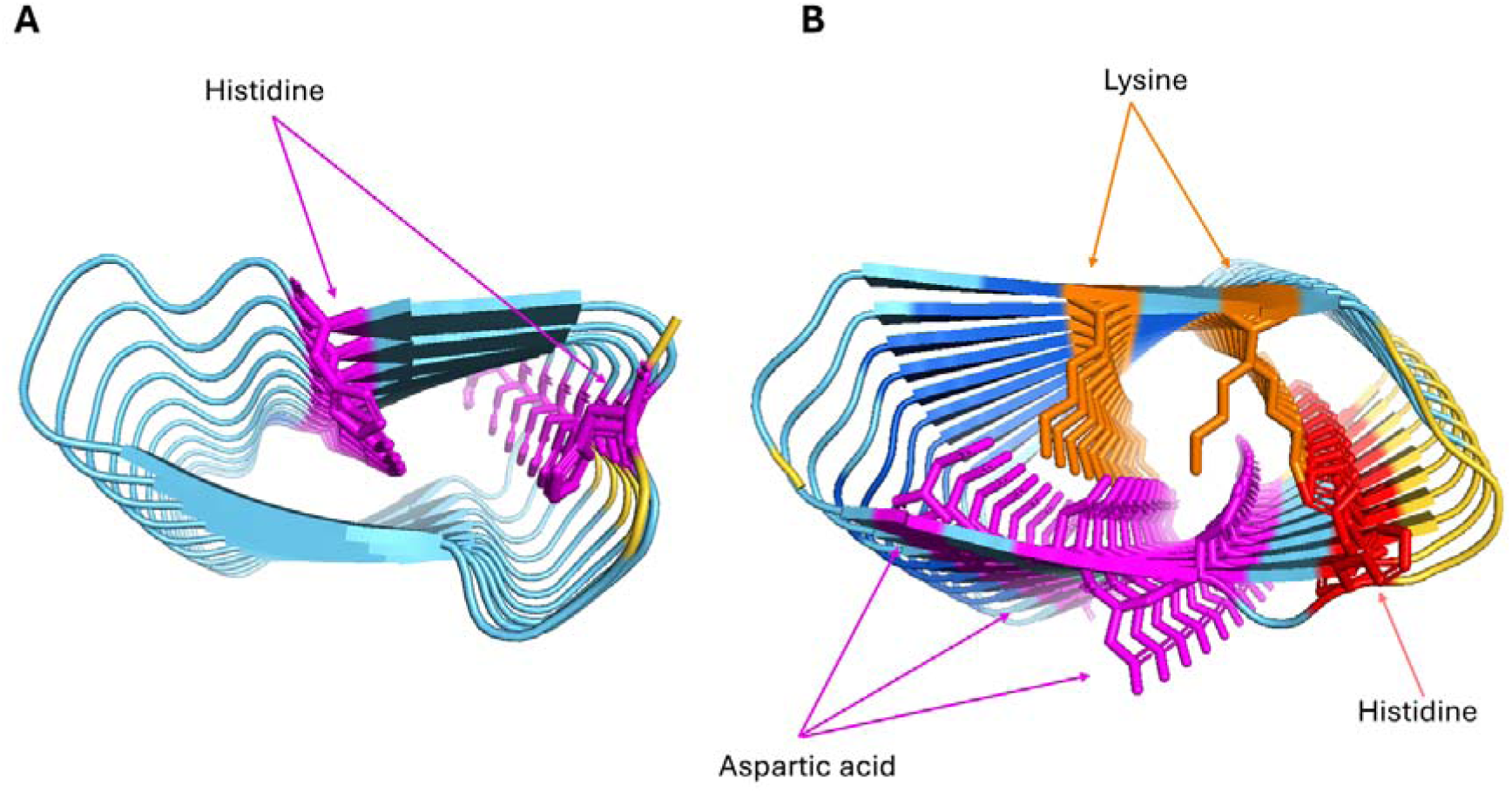
A) AF2 model with two internal histidine stacks despite a high pLDDT of 82.7. This model has a repeat unit length of 19 residues. B) AF2 model with several charged residues inside the protein despite the pLDDT of 82. In some cases, a salt-bridge may be formed, but the large amount of charge inside this protein is more likely to destabilise the structure. Significant clashing is also observed. This model has a repeat unit length of 21. Figure made using PyMOL (pymol.org).

### Alternative modelling methods show a stronger disagreement for ‘problematic’ structures

Alongside AF2, several other deep-learning protein prediction programs have been released in recent years. These include ESMFold, which uses a language model (Lin *et al*., 2023), and AF3, which incorporates a diffusion model (Abramson *et al*., 2024). As a comparison to AF2, all 32 sequences that gave a β-solenoid prediction with AF2 were modelled with both. The TM-scores between AF2 and ESMFold models and AF2 and AF3 models were then calculated for each sequence. Overall, both ESMFold and AF3 frequently disagreed with AF2 in structure, pLDDT or both (e.g., Figure 7).

**Figure 7:**
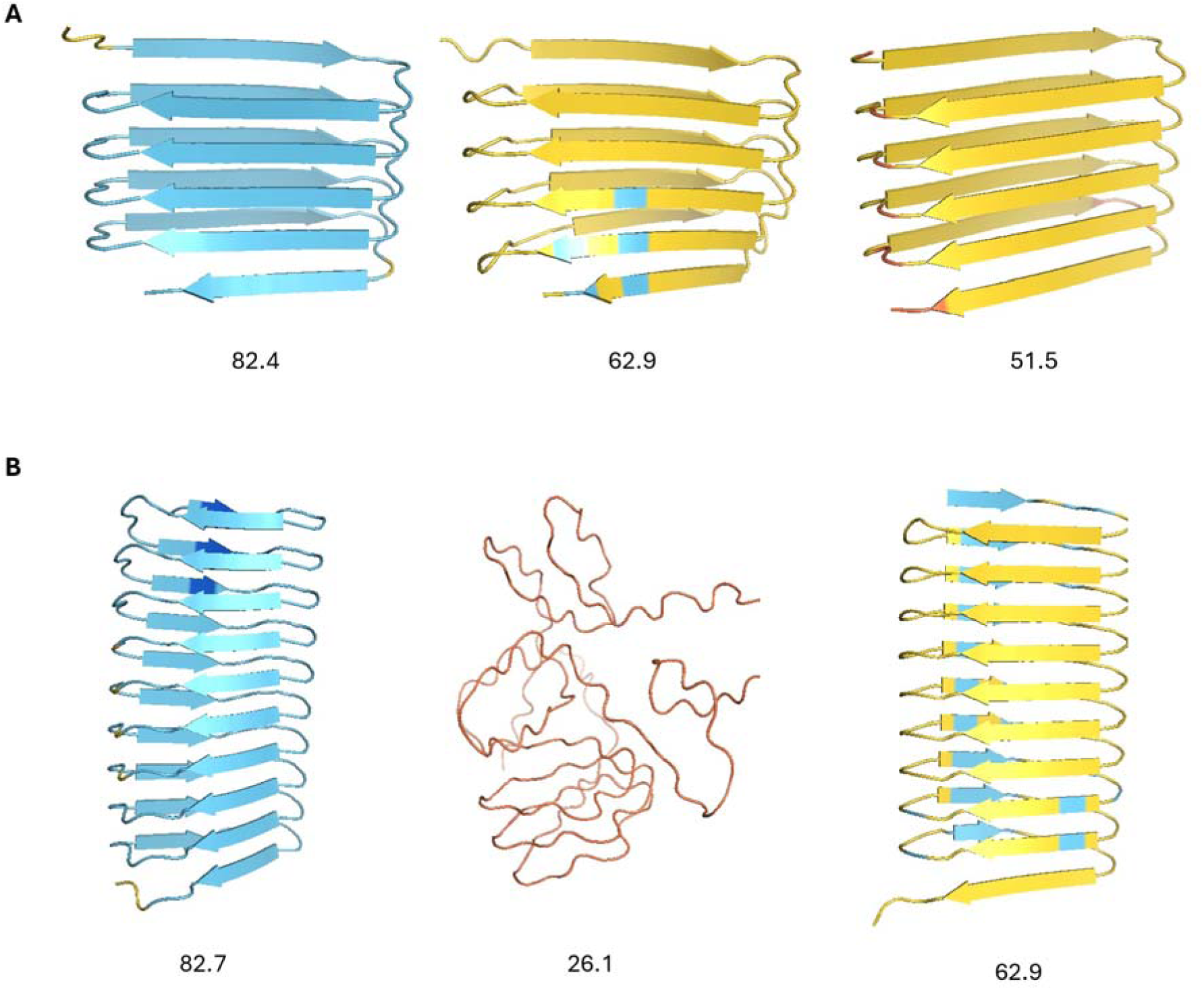
From left to right: AF2, ESMFold, AF3 model. pLDDT values are displayed beneath models. A) Example of a problematic model identified by Verify3D. B) Example of a problematic model with internally stacked histidine. All three methods disagree in pLDDT, despite sometimes producing similar structures. Figure made using PyMOL (pymol.org).

Figure 8 maps the structural divergence (difference in TM-score) between AF2 models and the predictions from ESMFold (x-axis) and AF3 (y-axis). Notably, a cluster of AF2 defined as ‘problematic’ are found near the origin. This set generally have much low pLDDT values (especially in the case of ESMFold) meaning that the sequences in question give high confidence β-solenoids with AF2, but very different and low-confidence predictions with the alternative methods. These results suggest that any blind spot AF2 may have with regard to perfect repeat sequences and β- solenoids, is not shared by ESMFold and AF3.

**Figure 8:**
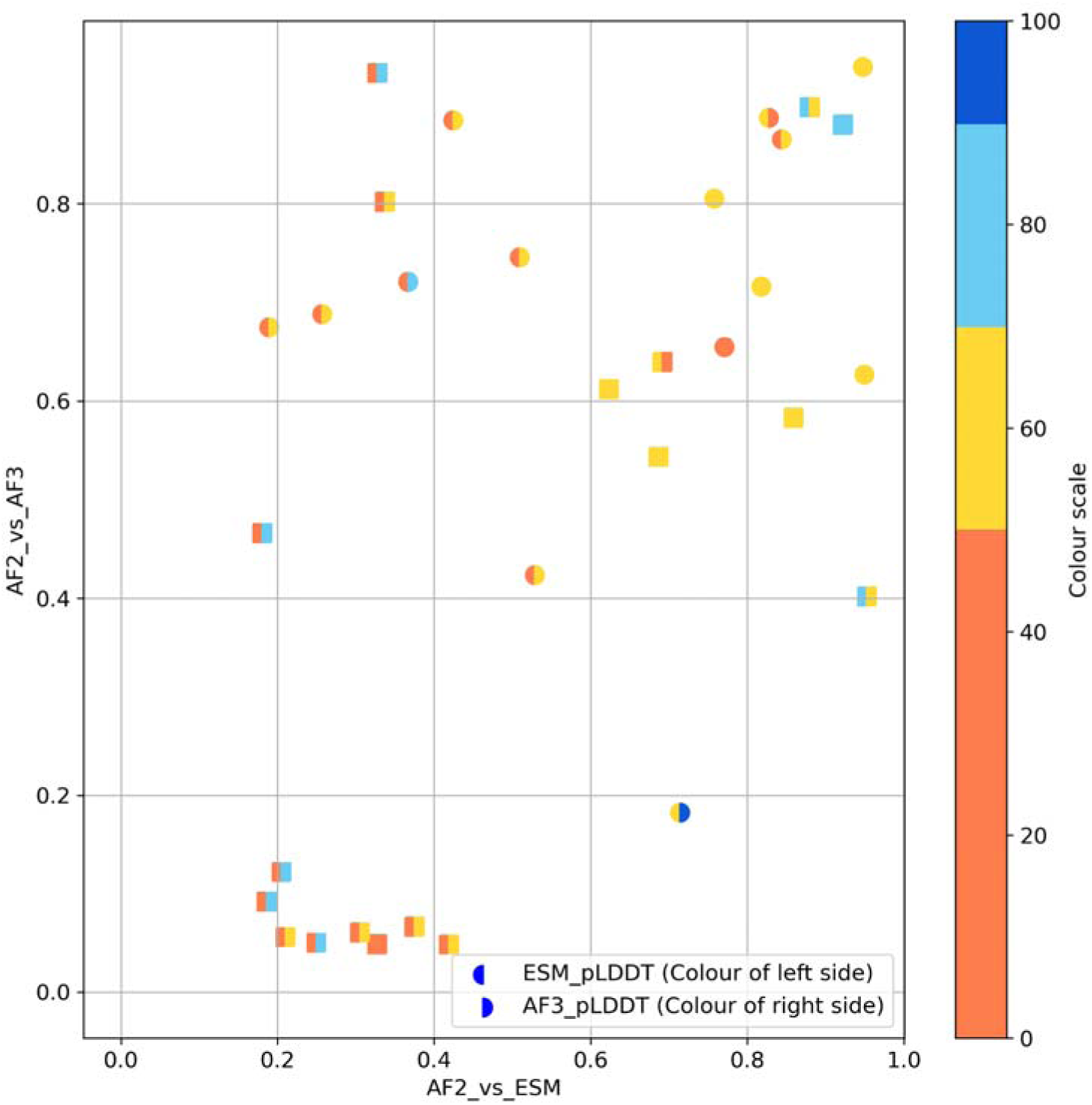
Comparison of AF2 models to ESMFold and AF3. The TM-score between AF2 and ESMFold models is shown on x-axis, and the TM-score between AF2 and AF3 is shown on the y-axis. Each point is coloured by both ESMFold pLDDT and AF3 pLDDT. Squares represent models that are considered to be ‘problematic’.

ESMFold also often predicted disordered structures. Additionally, AF3 sometimes generated long helical structures (e.g., Figure 9), consistent with the observation that it can ‘hallucinate’ such structure for sequence that is in fact intrinsically disordered (Abramson *et al*., 2024). This led to the hypothesis (tested below) that these proteins may actually be intrinsically disordered, despite AF2’s confidence in the β-solenoid fold.

**Figure 9:**
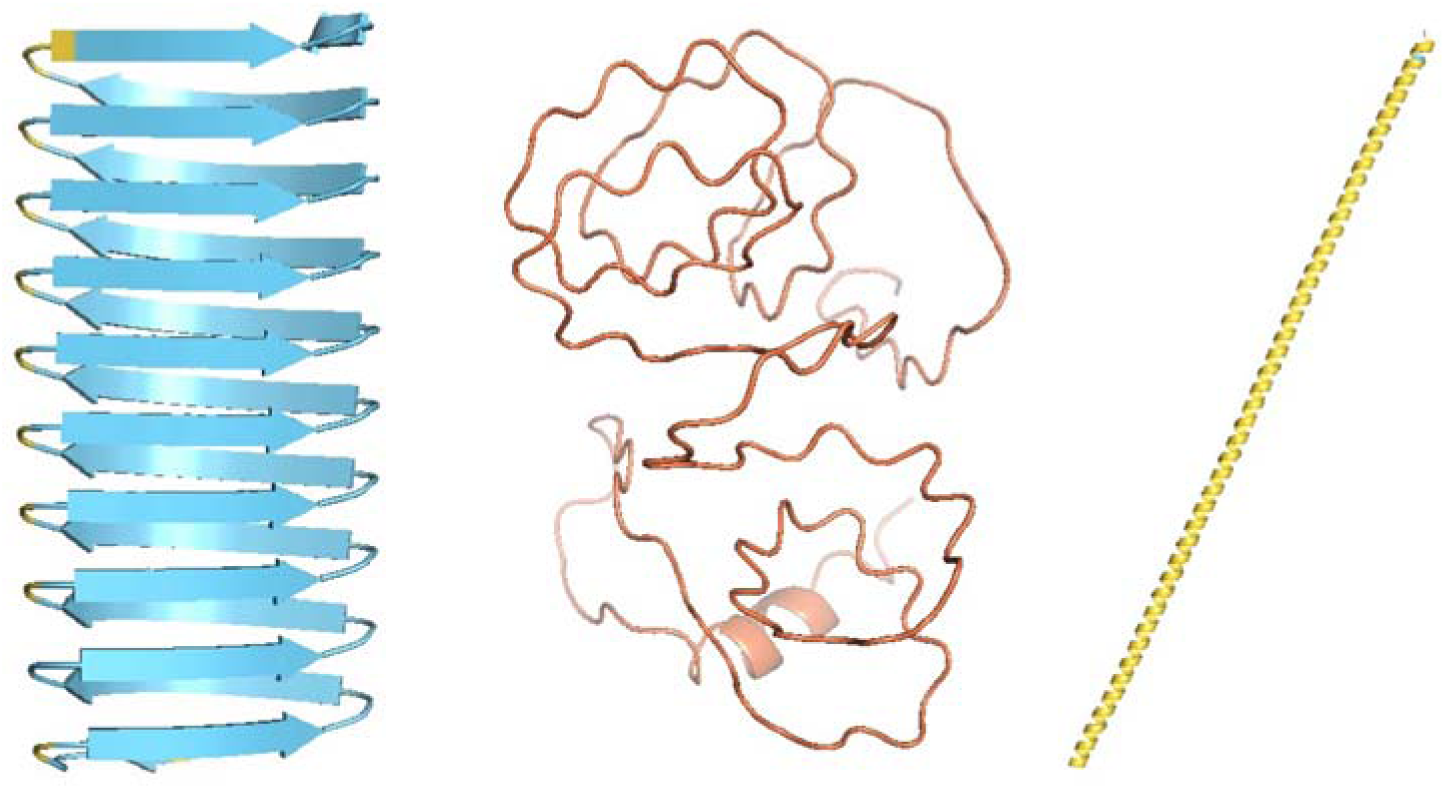
From left to right: AF2, ESMFold and AF3 for the same sequence (CITYVCWFYFYCFFCC). Models are coloured by pLDDT. It can be seen that ESMFold models this protein as disordered, despite the confident β-solenoid structure from AF2. AF3 displays what could be a helical ‘hallucination’ for an intrinsically disordered sequence. Figure made using PyMOL (pymol.org).

Finally, the sequences were modelled with RFAA (Krishna *et al*., 2024). For a large majority of the β-solenoid sequences, RFAA predicted a disordered structure (e.g., Figure 10), supporting the hypothesis that these proteins if synthesised would be disordered. To explore this idea, sequence disorder predictions with AIUPred were carried out (Erdős & Dosztányi, 2024).

**Figure 10:**
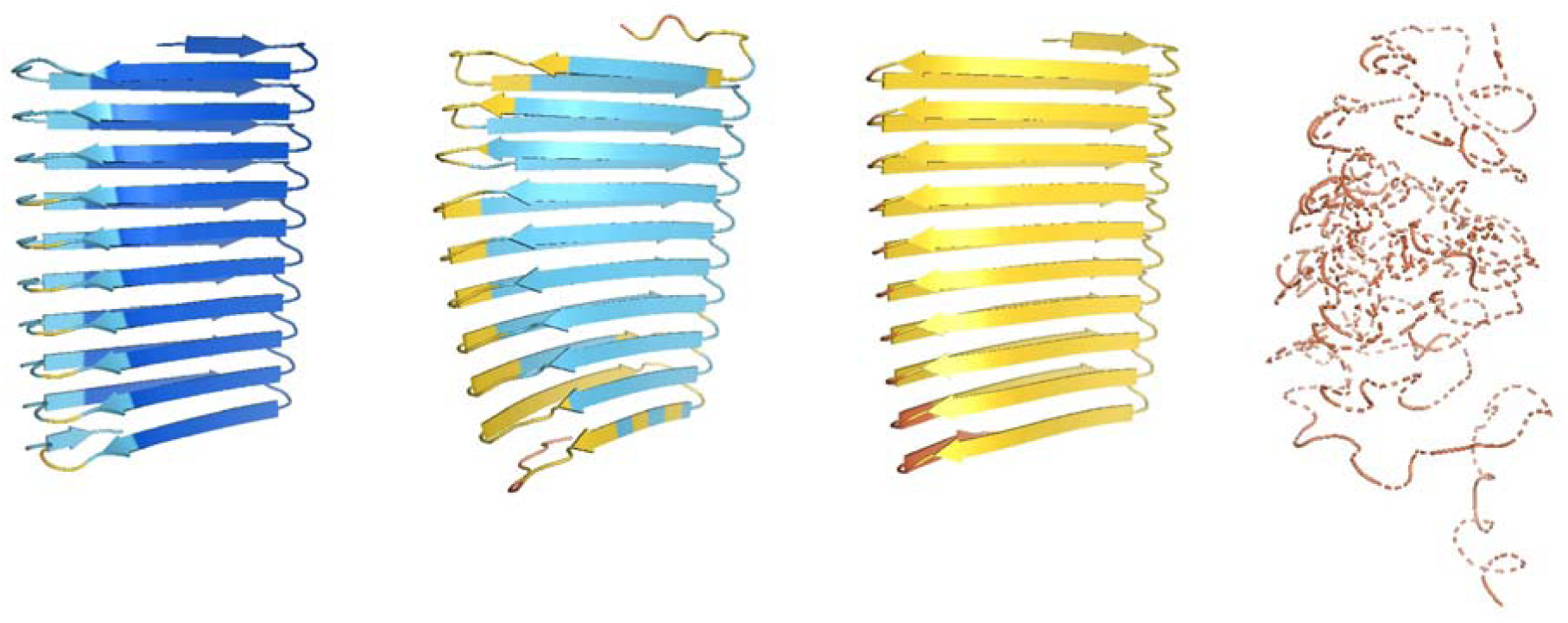
From left to right: AF2, ESMFold, AF3 and RFAA predictions for the same sequence (VCFCCFVTTCYITVWCYFYIFWCFI). An example of a confident β- solenoid model from AF2 showing as a disordered protein when modelled with RFAA. Figure made using PyMOL (pymol.org).

### Disorder predictions highlight another piece in the puzzle

Following disorder predictions, eight of the 32 β-solenoids had an average predicted disorder probability of ∼0.5 or above, seven of which had a pLDDT >70 (Figure 11). For those, ESMFold, RFAA, or both also predicted a disordered structure. This reinforces AF2’s bias surrounding β-solenoids whilst also supporting the idea that AF2 may have a tendency to confidently fold these disordered proteins into secondary structure due to their unusual repetition. Alternative modelling methods, however, seem less susceptible to this.

**Figure 11:**
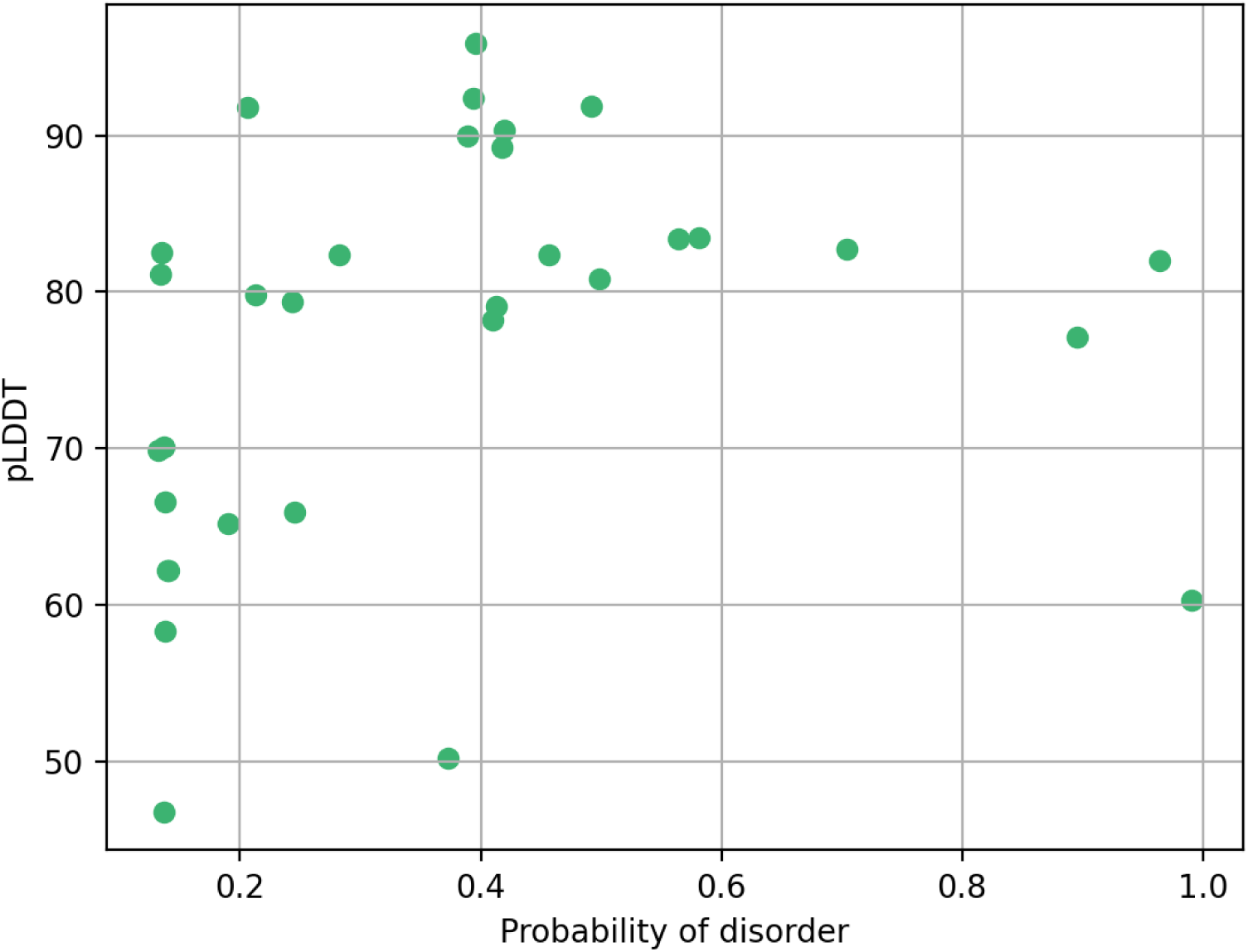
AIUPred disorder prediction results showing the pLDDT of each model on the y-axis and the average probability of disorder on the x-axis. Eight out of the 32 β- solenoid models were predicted likely to be disordered (probability ∼0.5 or above), seven of which have a pLDDT >70.

### HHPred search demonstrates occasional chance matching of artificial sequences to natural sequences

It was observed that RFAA rarely agreed with the other methods both structurally and in confidence. However, where there seemed to be a consensus across all methods, it was hypothesised that the artificial sequences may be matching to natural ones coincidentally, which could be influencing structure prediction. To investigate, the 32 sequences that AF2 modelled as β-solenoids were searched against the Pfam and PDB databases with HHPred (Zimmermann *et al*., 2018). Only three of the sequences produced significant hits, one of which was 6ZT4 from the Pentapeptide family (PF13599) (probability=98.4 , E-value <0.01) (Mistry *et al*., 2021). This family does consist of β-solenoid structures, which were included in the training data for the modelling methods. Figure 12 shows how this may have influenced predictions.

**Figure 12:**
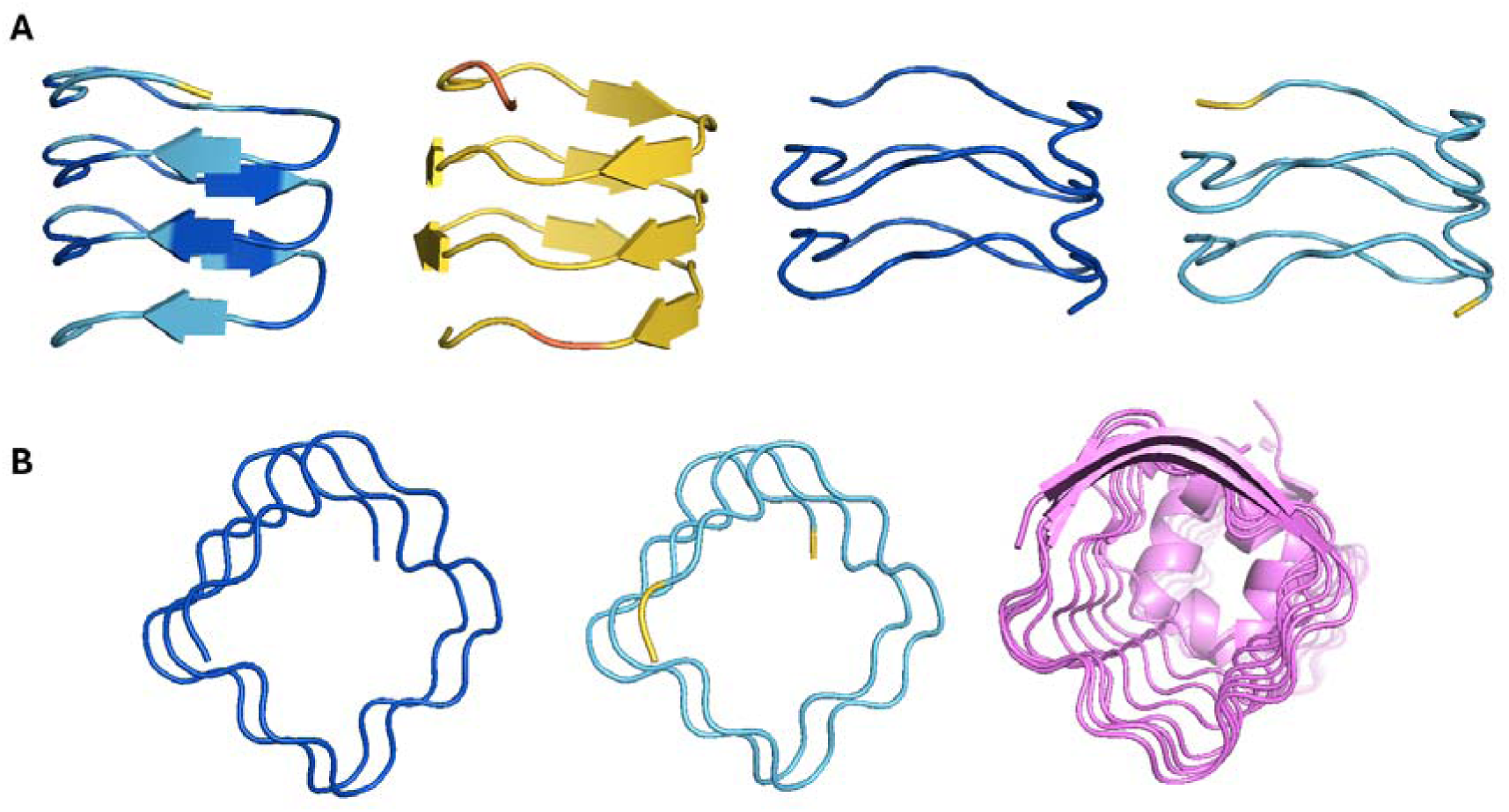
A) From left to right: AF2, ESMFold, AF3 and RFAA models for the same artificial repeat sequence (TGGDF) coloured by pLDDT. Both AF2 and ESMFold agree structurally, but not in pLDDT, and disagree with AF3 and RFAA, which do agree structurally and in confidence. B) From left to right: AF3 and RFAA models, and PDB structure 6ZT4. It appears that both AF3 and RFAA may have been influenced by this Pfam family (PF13599) by a chance matching of their sequences, explaining their structural similarity and high confidence. Figure made using PyMOL (pymol.org).

For some sequences, the false positive hits detected are likely due to overrepresentation of cysteine, a rare amino acid (Miseta & Csutora, 2000). It is possible that the different database search protocols implemented by the different Deep Learning-based methods are differently susceptible to detection of false positive hits. Figure 13 illustrates another example where HHpred found a chance similarity to PF19627 (Activity-dependent neuroprotector homeobox protein N-terminal) domain sequences that may have influenced the RFAA prediction but not the others.

**Figure 13:**
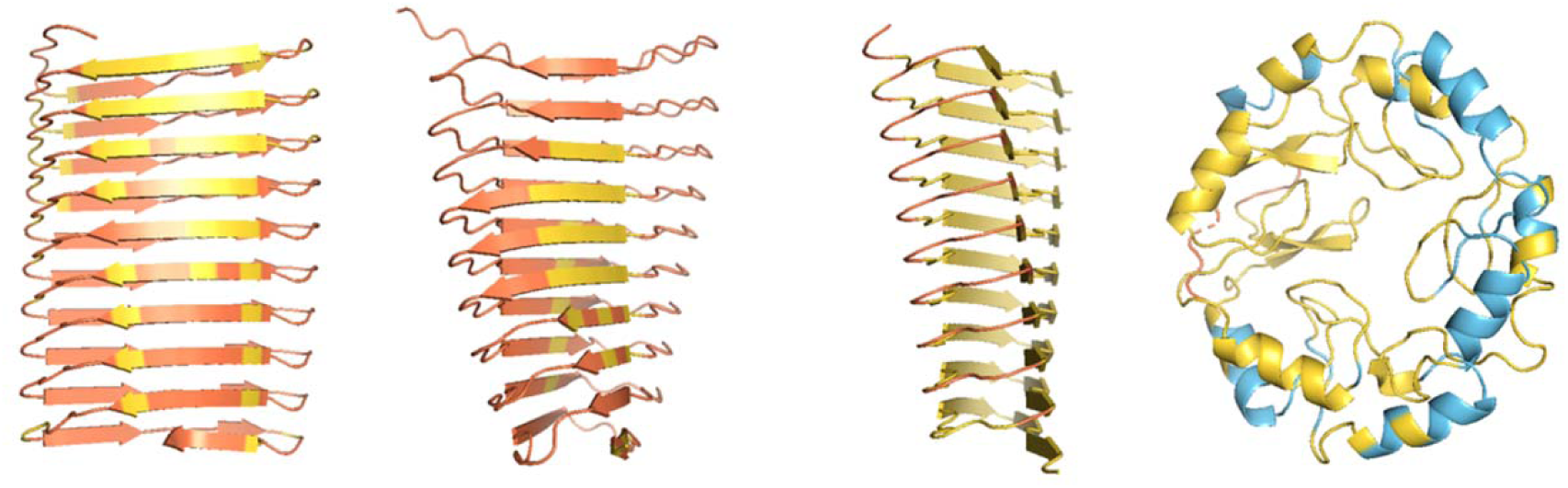
From left to right: AF2, ESMFold, AF3 and RFAA models for the same artificial repeat sequence (CMDIMPQHSYTCIPCQTFVPIEMNHR). It can be seen that RFAA disagrees structurally to the other methods, possibly reflecting differences in how each modelling tool implements its database search. Figure made using PyMOL (pymol.org).

### Molecular dynamics demonstrates model instability when compared to natural proteins

To test whether the models generated with high confidence, but internally stacked charged residues are unstable, all-atom MD was implemented with GROMACS.

Within the preliminary dataset, there were two high confidence models with internally charged stacks, both of which were the same size. These models were selected for simulation along with a natural β-solenoid crystal structure (PDB: 3JX8:A), in which there is no internal charged stacking.

The results of these simulations showed that the crystal structure remained stable throughout the simulation (Figure 14). In contrast, the models with negatively charged stacks were unstable. The model with internal glutamic acids (model 1 in Figure 14) became twisted during the simulation, albeit without unfolding completely: it may be that the observed aromatic stacks prevent further distortions. For the model with internal aspartic acids (model 2), the instability is even more evident with unfolding starting around the charged stack and leading to a distorted structure that is bent along the solenoid axis. The clear instability of these β-solenoid structures, recalling their high pLDDT values, again argues that AF2 has a blind sport with regard to some sequence repeat structures.

**Figure 14:**
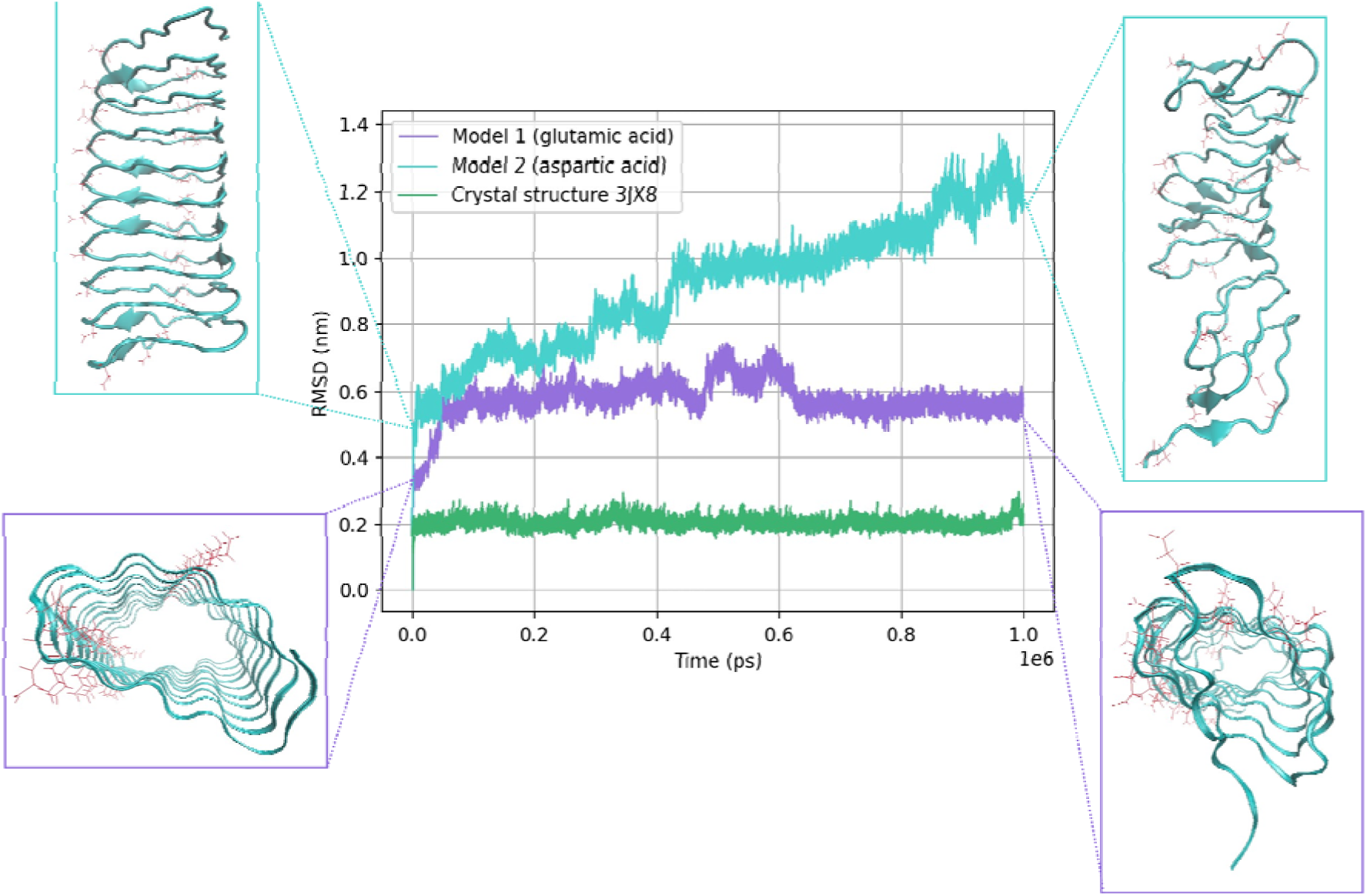
RMSD values over 1 microsecond for the three models tested with MD. Model structures are highlighted at the start and end of the simulation. Negatively charged residues are shown as red sticks. It can be seen that both models with internal charged stacks are more unstable than the crystal structure β-solenoid.

### β-solenoid bias may impact structure prediction of natural repeat proteins

As highlighted throughout this report, AF2 seems to have blind-spot surrounding β- solenoids when given random but ‘perfect’ repeat sequences. This has been demonstrated by the appearance of implausible features such as stacked charged residues, and quantified with Verify3D, which highlights model unfavourability despite AF2’s confidence. Additionally, when modelling the sequences with alternative modelling methods, a clear disagreement in confidence (and sometimes in fold prediction) reinforces this hypothesis. To test whether this bias towards β-solenoids is likely to influence prediction of natural proteins (some of which contain very near-perfect sequence repeats) we asked if the same pattern of higher pLDDT from AF2 compared to other methods was observed for these.

Accordingly, the sequences of Pfam families that AF2 had confidently predicted as possessing the β-solenoid fold (Mesdaghi *et al*., 2023), were modelled with the alternative deep-learning methods. For families PF07634, PF07012, PF02415, and PF01469, all four methods agreed structurally and in pLDDT, highlighting that these models are likely accurate. In each of these families the sequence repeats have diverged significantly (Table S1). Attention was then switched to near-perfect natural repeat sequences that may be influenced by this bias.

As Mesdaghi *et al*., (2023) show, AF2 predicts the human mucin proteins to adopt the β-solenoid fold with confidence. The sequence of mucin 1 in particular, closely resembles the artificial sequences in this study, consisting of an exact repetition of a 20-residue sequence twelve times. Despite this, AF2’s confidence is low. Mucin 22 on the other hand is less ‘perfect’, but AF2 predicts the β-solenoid fold with confidence. When modelling the whole protein in AF3, the confidence is lower, suggesting that this prediction may be problematic (Figure 15a, 15b).

**Figure 15:**
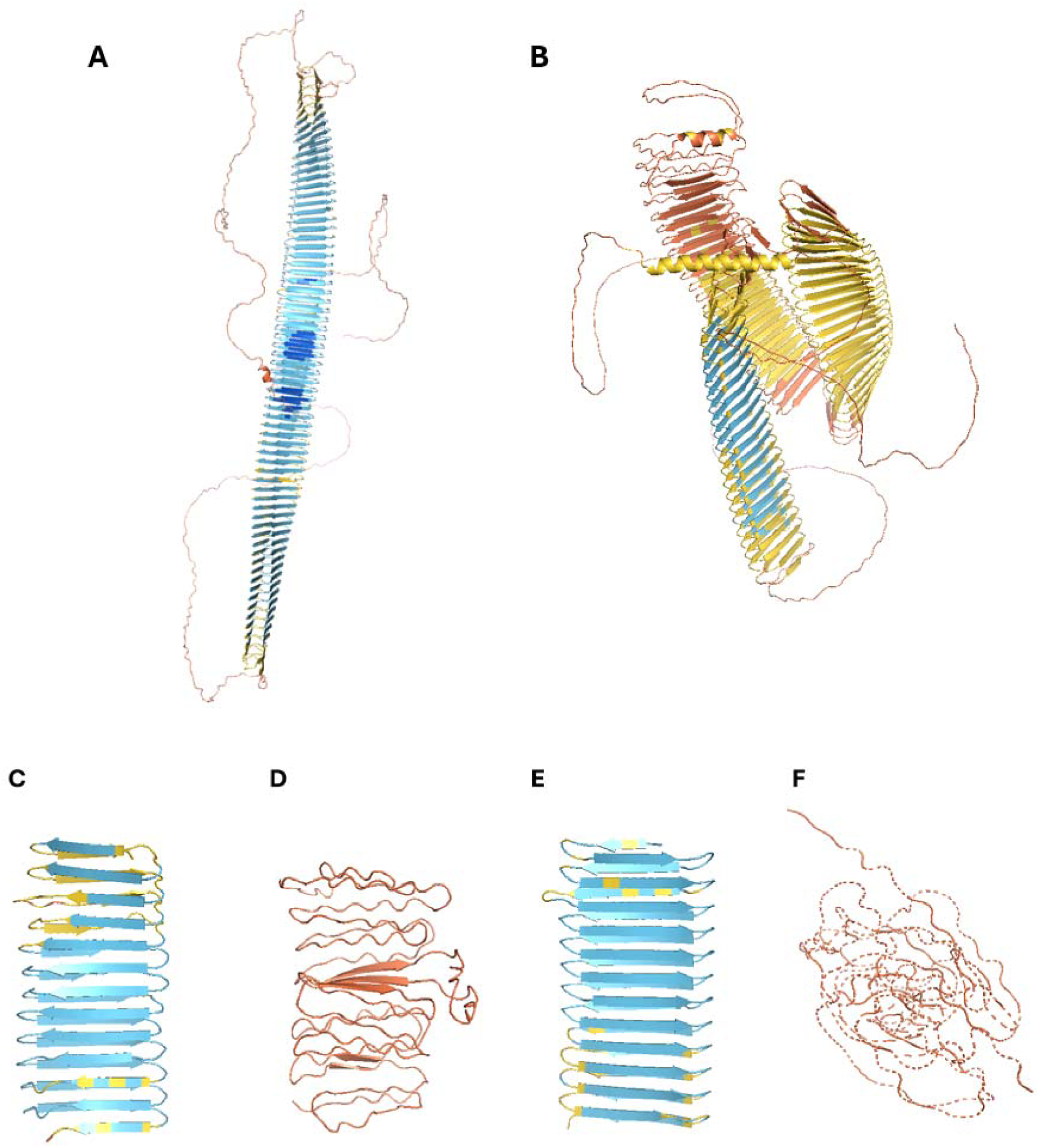
A) AF2 model of human protein mucin 22 from the AlphaFold Database (AFDB). B) AF3 model of mucin 22. C) AF2 model of mucin 22 fragment (residues 185-433). D) ESMFold model of mucin 22 fragment. E) AF3 model of mucin 22 fragment. F) RFAA model of mucin 22 fragment. Figure made using PyMOL (pymol.org).

Another pattern observed within this study was ESMFold and RFAA predicting disordered structures. Due to size constraints of ESMFold and RFAA, a fragment of mucin 22 (residues 185-433) was modelled for further comparison. Both AF2 and AF3 generated a β-solenoid model with confidence (Figure 15c, 15e); however, both ESMFold and RFAA predict a more disordered structure (Figure 15d, 15f). This suggests that this blind-spot has impacted this protein’s structure prediction, which should therefore be considered with scepticism.

## Conclusions & further studies

The emergence of AlphaFold 2 provoked an earthquake in protein structure prediction, offering the capacity to model most proteins to unprecedented accuracy. Nevertheless, as with any new tool, it is important to consider its strengths and weaknesses and recent work has described its relatively poor performance on predicting coiled-coil protein topology (Madaj *et al*., 2024) as well as a perceived blind-spot with regard to fold-switching proteins (Chakravarty *et al*., 2024). Evidence presented here suggests that AF2 has an unreasonable propensity to fold perfect sequence repeat proteins into β-solenoid structures, many of which – despite containing unusual and unstable features – it assigns high pLDDT values to.

The evidence for an AF2 blind-spot regarding β-solenoids is that more than 40% of random repeat sequences over a range of lengths fold that way, a large majority with average pLDDT >= 70 (moderate confidence) and several with pLDDT >90 (very high confidence). In some cases, there are inconsistences within AF2’s sets of five predictions – sequences that highly confidently predict as a β-solenoid in one model, but a similarly confident α-helix in another. And the models themselves contain features – external hydrophobic stacks and, particularly, internal charged stacks – that are rarely observed in experimental protein structures. These peculiarities are detected, to some extent, by the structure validation tool Verify3D which assesses protein structural viability by considering the secondary structure-aware environments different amino-acids. The internal charged stacks are demonstrated to lead to structural instability by Molecular Dynamics simulations. A quarter of sequences folded by AF2 into β-solenoids, all but one of moderate confidence or above, had mean intrinsic disorder scores from AIUPred of ∼0.5 or above suggesting that these sequences were in fact unlikely to have defined 3D folds. Finally, and importantly, alternative Deep Learning-based structure prediction tools, AF3, ESMFold and RFAA all behaved quite differently. Frequently, they produced quite different structure predictions but even when they did agree with the β-solenoid predicted by AF2 their confidence was much lower. Thus, only AF2 has the twin concern of implausible structure predictions combined with high confidence estimates.

These findings raise questions as to why AF2, but not other tools, exhibits this apparent blind spot. It is particularly interesting to compare the unreasonable internal stacking of charged residues in many models with the observation that AF2 is considered to have effectively learned an energy function (Roney & Ovchinnikov, 2022). Conceivably, the explanation lies, as mentioned, with the internal calcium binding sites observed in some β-solenoids (Garnham *et al*., 2011) at which the charge of acidic residues is compensated by the bound ion. Even though the models discussed here contain no positions resembling natural metal binding sites, AF2 might have learned that stacked acidic residues are acceptable in the context of the β-solenoid fold. Ultimately, the welcome and impressive ability of AF2 to accurately model ligand binding sites (Jumper *et al*., 2021), even though the ligands are absent from its calculations, may come at a price.

Another factor to consider is the potential for the randomly generated repeat sequences to accidentally resemble natural repeat proteins or known structures in general. The modelling methods used here differ in their approaches to capture known sequence-structure relationships, with AF3, for example, using a simpler procedure than AF2 to search for homologous sequences (Abramson *et al.,* 2024). However, chance resemblance to natural sequences and structures, revealed by HHPred analysis, appears to explain only a small part of the story: indeed, in the case of Pfam family PF13599 and its structure 6ZT4, only AF3 and RFAA models closely resembled the experimental structure.

Evidence from some natural repeat protein families suggests that the bias towards β- solenoids may only be consequential for repeat sequences that are (near-)perfect.

Future work could explore whether this bias affects more imperfect sequences. For example, for the sequences in this study, each rung of the solenoid could be mutated randomly to disrupt their ‘perfection’ and the same methods applied, although whether this would meaningfully mimic the process by which repeat proteins become less perfect, post-duplication is open to debate. It would also be interesting to follow up this work with perfect repeat sequences that better followed natural amino-acid abundance rather than, as here, representing each amino-acid with equal frequency. Ultimately, experimental validation of β-solenoids predicted for natural proteins will be required for any confidence that the AF2 blind spot around β-solenoids affects only theoretical, perfect repeats.

Despite the fully justified excitement around AF2 and following structure prediction tools, it remains important to fully understand their strengths, weakness and quirks; particularly so since, like many Deep Learning-based tools, they have limited interpretability (Tan & Zhang, 2023). Exercises focused on performance against specific fold types, especially those that might be under-represented in training sets, are therefore important for each new tool that emerges (eg Wojciechowska *et al.,* 2024). As computed structures (Varadi *et al.,* 2024; Steinegger *et al*., 2024) become as broadly available as experimental structures, this validation is more important than ever: only when all limitations and biases are comprehensively understood, can computational methods illuminate all uncharted areas of the protein universe.

