## Supplementary Material for "AlphaFold 2, but not AlphaFold 3, predicts confident but unrealistic β-solenoid structures for repeat proteins"

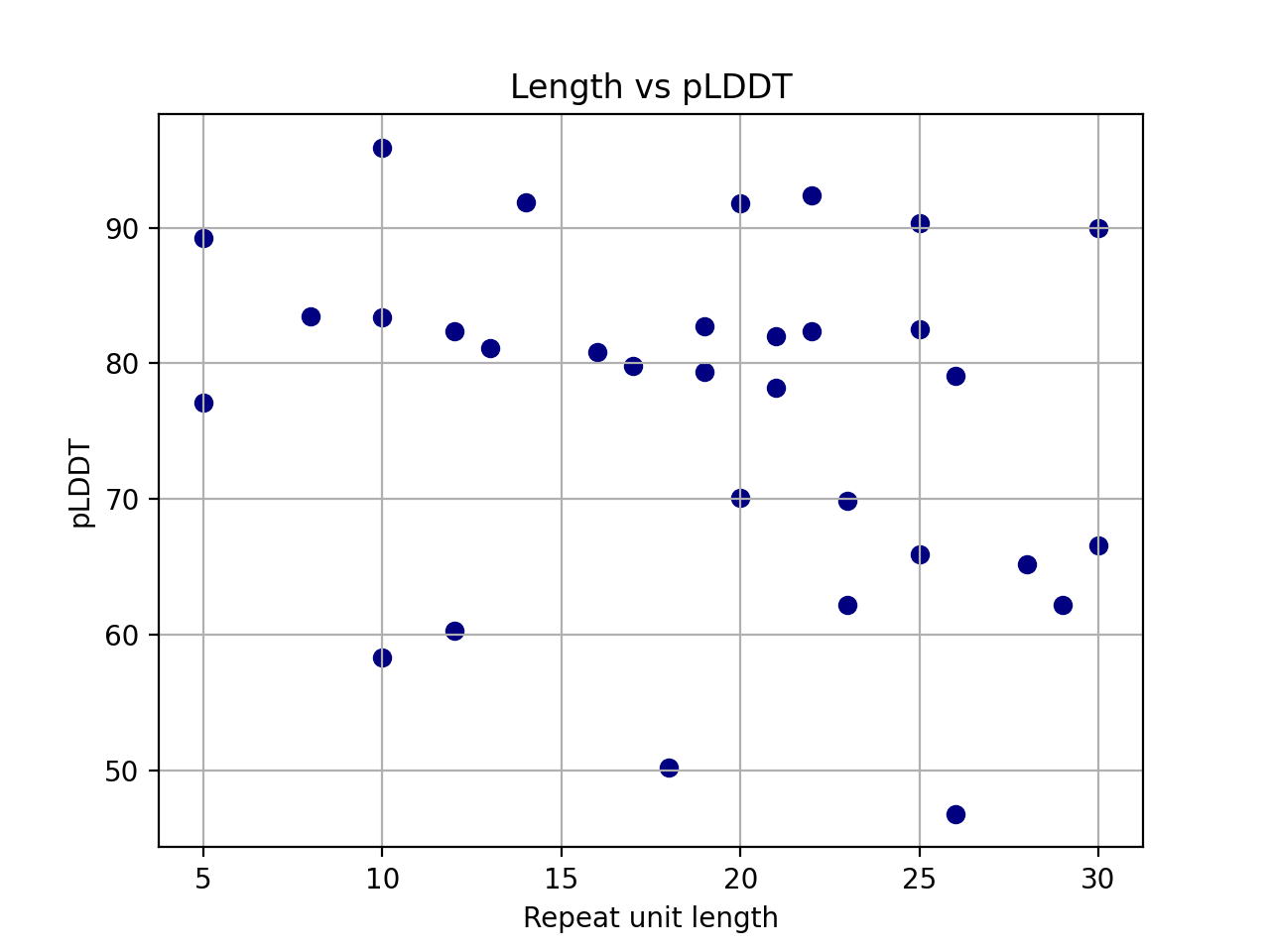

***Figure S1:*** *The repeat unit length of each β-solenoid model plotted against the pLDDT. There are high pLDDT models (>70) across all repeat unit lengths.*

***Table S1:*** *The representative Pfam family sequence repeats as detected by RADAR showing significant sequence divergence post-duplication.*

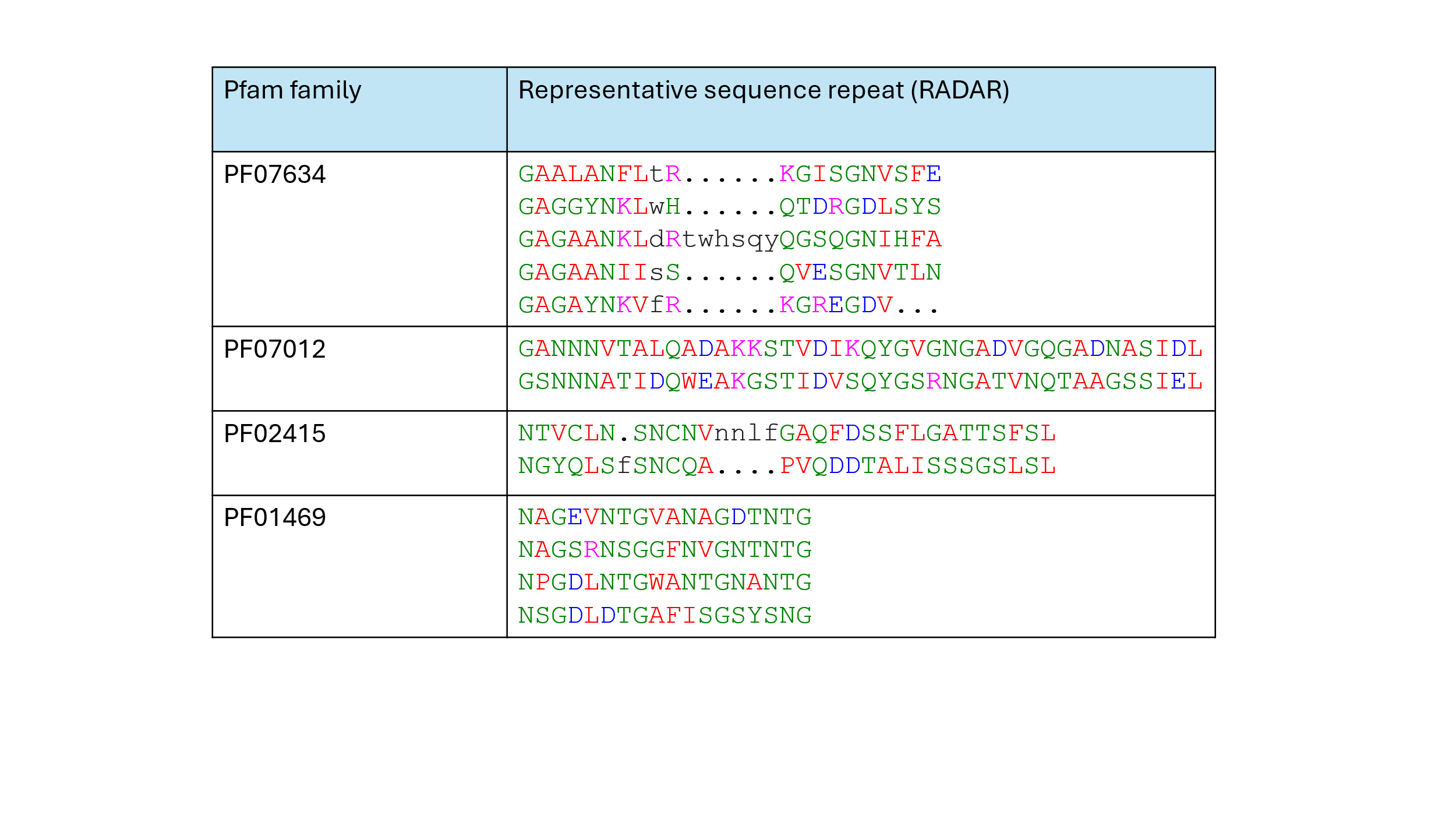
